# *The Binary Cellular Biology of Human Growth*: II. Calculating the Size of the Human Fetus from the Size of its Parts

**DOI:** 10.1101/2025.01.22.634333

**Authors:** Zifan Gu, Rashi Gupta, David E Cantonwine, Thomas F McElrath, Henning Tiemeier, James Selib Michaelson

## Abstract

Assessing and understanding the ***Size*** of the human fetus provides essential information for the management of the health of the newborn and its mother, a quality that is commonly measured by ***Fetal Ultrasound***. Many equations for making such calculations have been considered, none of which have been entirely satisfactory. As we have shown in the previous paper in this series, a consideration of the formation of the body, in units of numbers of cells, ***N***, an approach we call ***Binary Cellular Analysis***, provides new quantitative tools for estimating the ***Size*** and ***Age*** of the fetus. These tools include a new mathematical basis for creating useful expressions, ***Binary Cellular Estimated Fetal Weight Equations***, for calculating human ***Fetal Weight*** from ***Ultrasound Measurements***. As we show here, the most promising of these expressions, the ***Abdominal Circumference Binary Cellular Estimated Fetal Weight Equation***, performs better than other currently used ***Fetal Weight Equations***, while also yielding up new methods for improving ultrasound size assessment. Web-based calculators (https://kidzgrowth.com) make these mathematical manipulations available for obstetric care.

## INTRODUCTION

While many equations have been considered for estimating ***Fetal Weight*** from ***Ultrasound Measurements***, none have been entirely satisfactory^.1-14^ To address this challenge, in this, the second of three articles in this series,^15,16^ we again return to a consideration the ***Size*** the whole body, ***w***, and its parts, ***p***, in units of numbers of cells, ***N***_***w***_ and ***N***_***p***_, especially those parts measured by ***Fetal Ultrasound***. We call this method ***Binary Cellular Analysis***, as it examines the ***Binary*** choices made by our cells to be ***Mitotic*** or ***Quiescent***, determining the number of cells in our ***Bodies*** and our ***Parts***.^17-19^

As we have seen in the first article in this series,^16^ a ***Binary Cellular*** approach provides new expressions for calculating the ***Size*** and ***Growth*** of human fetuses from ***Fetal Ultrasound Measurements: Binary Cellular Estimated Fetal Weight Equations***.^16^ Here we shall see that the most promising of these expressions, the ***Abdominal Circumference Binary Cellular Estimated Fetal Weight Equation***, performs better than other widely used ***Estimated Fetal Weight Equations***, while also yielding up new methods for improving ultrasound size assessment.

## METHODS

### The *Combined Dataset*

For this, and the following manuscript in this series,^15^ we utilized a deidentified dataset of 821 pregnant women seen at the Brigham and Women’s Hospital. We call this dataset the “***Combined Dataset***”. In addition to ***Birthweight***, these datasets contained information on ultrasound measurement of: ***Biparietal Diameter*** (BPD), ***Occipitofrontal Diameter*** (OFD), ***Abdominal Diameter*** (AD), ***Femur Length*** (FL), ***Head Circumference*** (HC), ***Abdominal Circumference*** (AC), and ***Crown Rump Length*** (CRL).

The ***Combined Dataset*** was divided randomly into to two subsets: a ***Training Dataset***, used to derive the parameters for the new ***Binary Cellular Estimated Fetal Weight Equations***, and a ***Testing Dataset***, used to test the accuracy of these expressions and parameters. Both datasets contained 117 ultrasound examinations carried out within 7 days of birth.

Here we report the general findings of our analysis of the estimation of fetal weight from ultrasound. For an exhaustive description of the underling calculations, see the SUPPLEMENT appended to this communication.

Data were analyzed in Microsoft Excel.

### Terminology for *Fetal Growth*: *Age, Gestational Age, Conceptional Age, Fetal Age*

***Gestational Age***, the time since the last menstrual period, has long been the most common way for thinking about the age of the human fetus. Another commonly used term is ***Conceptional Age***, but the date of conception is not always known. As we are examining fetal, childhood and adult growth data, as well as growth data of animals such as mollusks, tunicates, and nematodes, which don’t have a fetal period, we shall be using the simplest term, ***Age***, with modifiers to indicate the period examined, which, for our work here, will be ***Fetal Age***. Here we shall calculate ***Fetal Age*** from ***Crown to Rump Length*** (***CRL***) ***Ultrasound*** measurements.

### Equations not derived or specified in the main text of this communication

As developed in the previous paper in this series,^16^ the ***Binary Cellular Universal Growth Equation*** (#2), for capturing the relationship between ***size, N***, and ***age, t***:

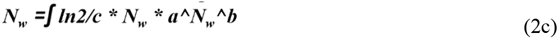

To calculate human ***Fetal Age***, we have adopted and adapted the INTERGROWTH Equation for estimating ***Fetal Age*** from ***Crown to Rump Length*** (***CRL***)^20^:

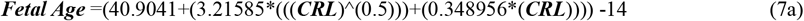

We call this expression, the ***CRL Fetal Age*** ***Equation*** (#7a).^16^ The 14 days subtracted from the INTERGROWTH value converts the values from the ***Gestational Age*** to ***Fetal Age***.

## RESULTS

### The Data: *Ultrasound Measurements* vs *Birthweight*

#### *Ultrasound Measurements* vs *Birthweight*: The Relationship

Let us begin by taking a general look at the relationships between various ***Ultrasound Measurements*** and ***Birthweights*** in our datasets (METHODS section). This can be seen simply by graphing ***Ultrasound Measurements*** made shortly before birth against the corresponding ***Birthweights*** and fitting the data to a simple linear regression. Such graphs for ***Ultrasound Measurements*** carried out within 7 days of birth can be seen in **Figure 1** and **Supplement-Figure-II.1**. These graphs reveal that for each kilogram of ***Fetal Weight, Abdominal Circumference*** (AC) increases by ∼447 millimeters, while ***Abdominal Diameter*** (AD) increases by ∼142 millimeters, ***Head Circumference*** (HC) by ∼19.6 millimeters, ***Occipitofrontal Diameter*** (OFD) by ∼7.8 millimeters, ***Biparietal Diameter*** (BPD) by ∼6.2 millimeters, and ***Femur Length*** (FL) by ∼5.9 millimeters (**Figure 1** and **Supplement-Figures II.1&2**).

**Figure 1.**
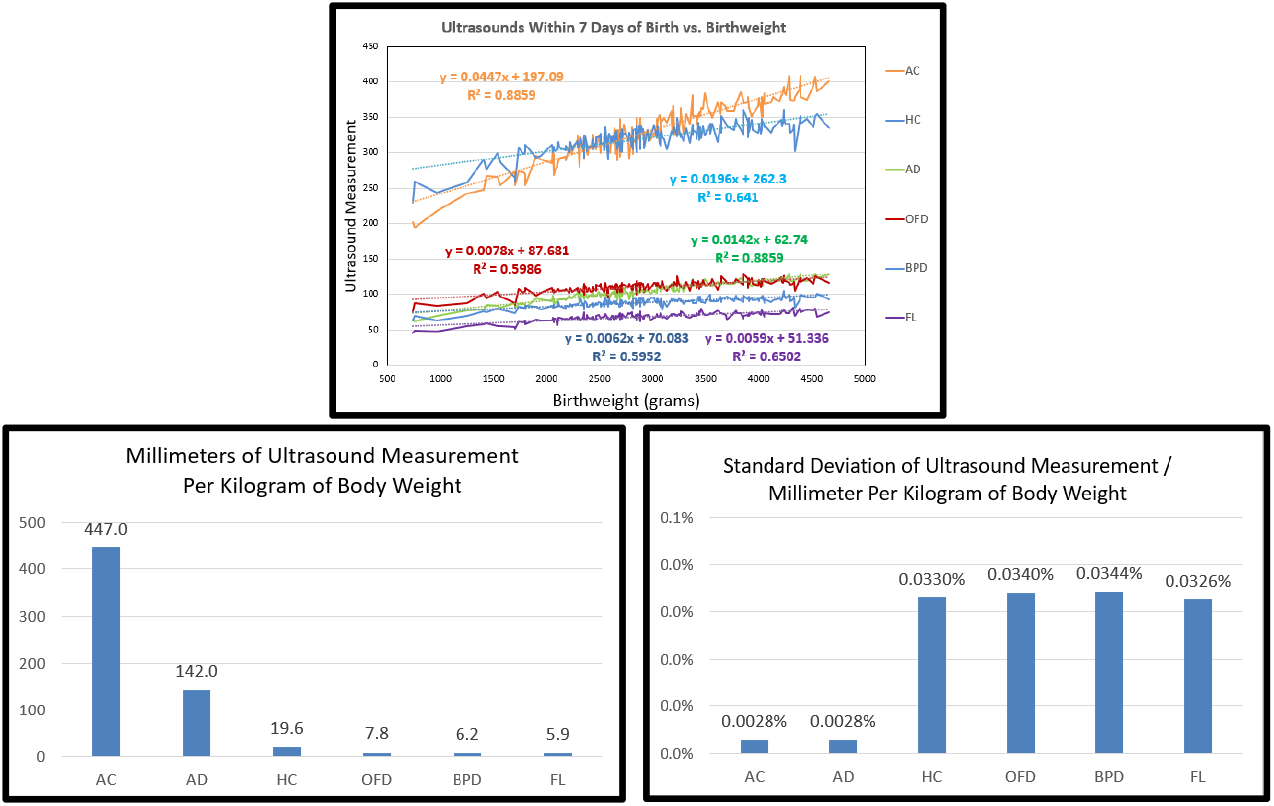
Comparison of ***Birthweights***, and ***Ultrasound Measurements*** made within 7 days of birth. TOP: Fit of the data to linear regressions. MIDDLE: Millimeters/Kilogram. BOTTOM: Standard Deviation.

#### *Ultrasound Measurements* vs *Birthweight*: The Noise

We can visualize the noise of each ***Ultrasound Measurement*** simply by constructing scatter plots of ***Birthweight*** vs. ***Ultrasound Measurements*** made within a few days of birth. Note that the ***r***^***2***^ value for ***Abdominal Measurement*** is much smaller than for other anatomical ***Measurements*** (**Supplement-Figure-II.3**). The ***Abdominal Circumference*** (AC) measurements also had a much lower ***Standard Deviation*** than the other ***Ultrasound Measurements***. (**Figure 1** and **Supplement-Figure-II.2**).

#### *Ultrasound Measurements* vs *Birthweight*: The Relationship Vs The Noise

This first look at the relationship between ***Ultrasound Measurement*** and ***Birthweight*** reveals that ***Abdominal Circumference*** (AC) gives much greater leverage in capturing ***Fetal Weight*** than the other ***Ultrasound Measurements***. For example, ***Abdominal Circumference*** (AC) had 75-fold greater potential to capture change in ***Fetal Weight*** than ***Femur Length*** (FL), with 10-fold less error. Thus, ***Abdominal Circumference*** (AC) gives a much stronger, and much less noisy, signal for estimating ***Fetal Weight*** than the other ***Ultrasound Measurements***. These features make clear why ***Abdominal Circumference*** (AC) has repeatedly been proven to be the most effective ***Ultrasound Measurement*** for many ***Estimated Fetal Weight Equations***. The biological reasons for this will be outlined in the DISCUSSION.

#### Current *Estimated Fetal Weight Equations* for Determining *Fetal Weight* from *Ultrasound*

Many ***Estimated Fetal Weight Equations*** have been constructed for estimating the size of the fetus from ultrasound measurements.^1-14^ Among the most common are:

***Hadlock Estimated Fetal Weight Equation-1***:

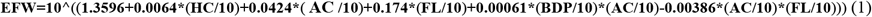

***Hadlock Estimated Fetal Weight Equation-2***:

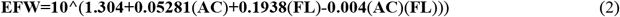

***Hadlock Estimated Fetal Weight Equation-3:***

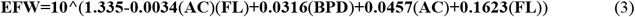

***Hadlock Estimated Fetal Weight Equation-4***:

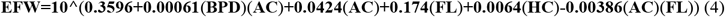

***Hadlock Estimated Fetal Weight Equation-5***:

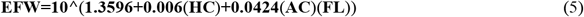

***INTERGROWTH Estimated Fetal Weight Equation***:

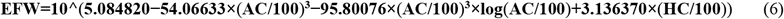

***Hadlock Estimated Fetal Weight Equation-4*** is the most commonly used ***Estimated Fetal Weight Equation***, and many studies have found it to be superior to the others.^1-14^

#### New *Binary Cellular Estimated Fetal Weight Equations* for Determining *Fetal Weight* from *Ultrasound*

As we described in the previous paper in this series,^16^ the analysis of the ***Size*** of many parts of the body (***tissues, organs***, and ***anatomical structures***), in units of numbers of cells, ***N***_***p***_, when compared with the ***Size*** of the body as a whole, ***N***_***w***_, usually shows the relationship:

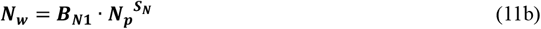

We call this expression the ***Binary Cellular Allometric Growth Equation***, and the form, and parameters, of this expression capture how the body makes its parts from its cells. As we shall outline in the DISCUSSION, the cellular nature of this expression gives the biological reasons for why some ultrasound measurements give better information on the ***Size*** of the body than other ultrasound measurements.

As also described in the previous paper in this series,^16^ with this expression, we could put ***Binary Cellular Biology*** to work to derive a new, biologically based, method for calculating ***Fetal Weight*** from ***Ultrasound Measurements, Binary Cellular Estimated Fetal Weight Equations***:

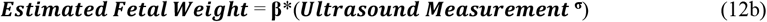

We call this expression the ***1st-Step-Binary Cellular Estimated Fetal Weight Equation***.

A problem that has arisen for many ***Estimated Fetal Weight Equations*** has been the inability to accurately capture the sizes of fetuses that will become ***Large*** and ***Small Babies***^.27,31^,***36 This imbalance is easily solved by adding one more parameter, γ***:

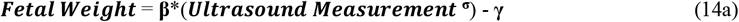

or

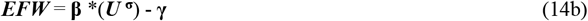

*We call Equation #14 the* ***Binary Cellular Estimated Fetal Weight Equation***.

Identifying the values for the **β, *σ***, and **γ** parameters is tedious, but straightforward, and the process of doing so with data from the ***Training Dataset*** (see METHODS) is described in detail in the SUPPLEMENT. For ***Abdominal Circumference***: **β=*0*.*001371156***; ***σ=2*.*5187***; **γ=*244*.*25*** and thus:

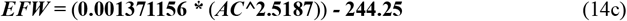

We call Equation #14c the ***Abdominal Circumference Binary Cellular Estimated Fetal Weight Equation***.

#### Testing *Estimated Fetal Weight Equations*

##### Overview

Having determined the values for the **β, *σ***, and **γ** parameters of the ***Binary Cellular Estimated Fetal Weight Equations***, derived from the ***Training Dataset***, we then tested their performance against the ***Testing Dataset***, along with other widely used ***Estimated Fetal Weight Equations*** (Equations #1-6). These tests were exhaustive, and interested readers can find these details in the SUPPLEMENT.

##### *Mean* and *Median Error*

The ***Mean*** and ***Median Errors*** and ***Absolute Errors*** for the ***Abdominal Circumference Binary Cellular Estimated Fetal Weight Equation*** (#14c) proved to be superior to these generated by the ***INTERGROWTH*** and ***Hadlock-1,-2,-3,-4***, and ***-5 Estimated Fetal Weight Equations*** (**Figure 2** and **Supplement-Figures-II.9** and **10**). For the ***Abdominal Circumference Binary Cellular Estimated Fetal Weight Equation*** (#14c), the ***Mean Error*** was 0.5%, while the ***Hadlock-4 Estimated Fetal Weight Equation*** had a ***Mean Error*** of 5.4%, with the ***INTERGROWTH*** and ***Hadlock-1,-2,-3*** and ***-5 Estimated Fetal Weight Equations*** having higher values.

**Figure 2.**
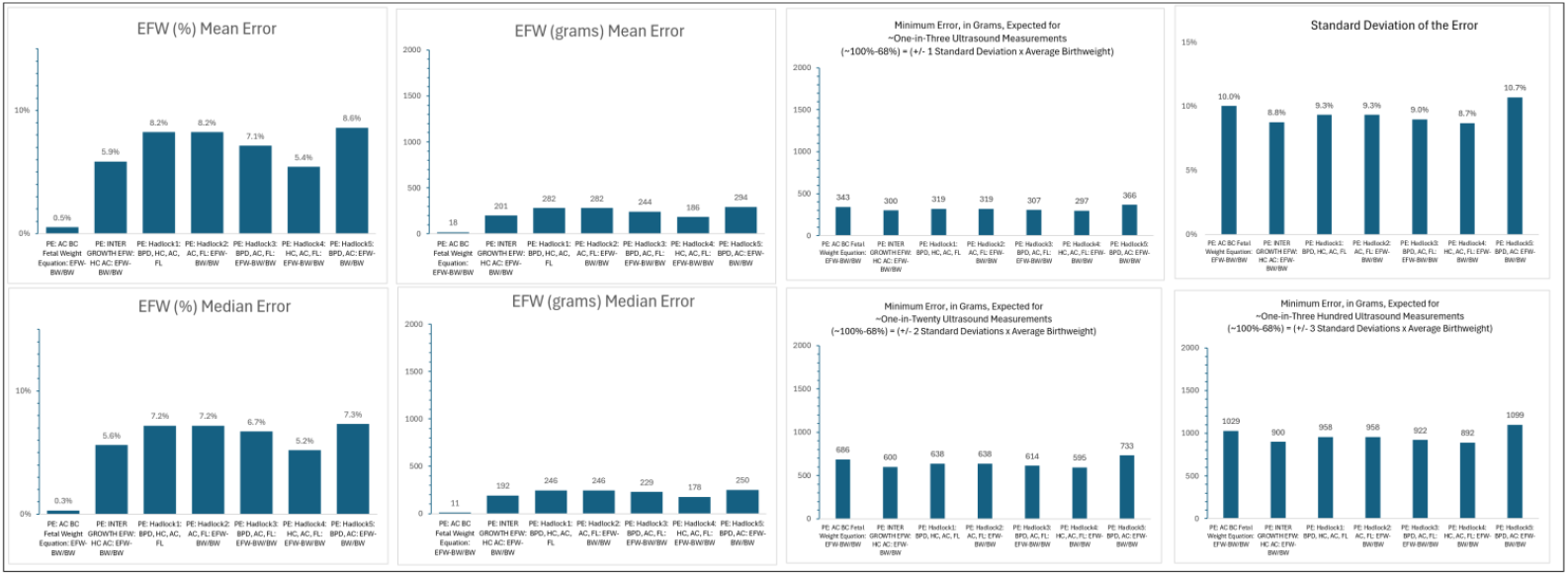
***Mean*** and ***Median Errors*** and ***Standard Deviation*** of the ***Error***, of seven ***Estimated Fetal Weight Equations: Abdominal Circumference Binary Cellular Estimated Fetal Weight Equation, INTERGROWTH Estimated Fetal Weight Equation, Hadlock-1,-2,-3,-4***, and ***-5 Estimated Fetal Weight Equations***, in the ***Testing Dataset***, based on ultrasound carried out within 7 days of birth.

##### Standard Deviation

The ***Standard Deviations*** of all seven ***Estimated Fetal Weight Equations*** were similar. The ***Hadlock-4 Equation***, at 8.7%, was very slightly better than the ***Abdominal Circumference Binary Cellular Estimated Fetal Weight Equation*** (#14c), at 10.0% (**Figure 2** and **Supplement-Figure-II.9**). However, the practical difference is negligible. For ∼1-in-300 ultrasound measurements, the difference between the ***Abdominal Circumference Binary Cellular Estimated Fetal Weight Equation*** (#14c), and the ***Hadlock-4 Estimated Fetal Weight Equation***, was a modest 137 grams, roughly 4% of the weight of an average newborn. For 2 out of 3 ultrasound measurements, the difference between was an even more modest, at 28 grams, roughly 1% of the weight of an average newborn. The ***Standard Deviations*** of the ***INTERGROWTH*** and ***Hadlock-1,-2,-3*** and ***-5 Estimated Fetal Weight Equations*** had slightly higher values than the ***Hadlock-4*** and ***Abdominal Circumference Binary Cellular Estimated Fetal Weight Equation*** (#14c).

The Problem isn’t the ***Error***, but the ***Standard Deviation***, Caused by the Sonographers, Not the Equations. While there was little difference in the ***Standard Deviations*** of all seven ***Estimated Fetal Weight Equations***, for all seven, these ***Standard Deviations*** were disturbingly large. As we shall see below, the source of this degradation of ***Estimated Fetal Weight*** lies not in the equation used, but in the performance of the ultrasound measurements by the sonographer.

##### The Difference Between *Error* and *Standard Deviation*

Simply put, a calculation of the ***Error*** tells us how much estimates of ***Fetal Weight*** are off for ***all of the fetuses, on average***, while a calculation of the ***Standard Deviation*** tells us how much estimates of ***Fetal Weight*** are off for ***the few of the fetuses*** that are ***on the extreme***. The ***Error*** tells us ***Where*** the ***PEAK*** of the ***Bell Curve*** is. The ***Standard Deviation*** tells us ***How Wide*** the ***FLANKS*** of the ***Bell Curve*** are. The calculations of ***Estimated Fetal Weight*** shown here tell us that ***PEAK*** of the ***Bell Curve*** is roughly where it should be, but the ***FLANKS*** of the ***Bell Curve*** stretch out too far (**Figure 2** and **Supplement-Figures-II.9&10**).

##### The Difference Between *Estimated Fetal Weight Error* and *Standard Deviation*

Among all 7 ***Estimated Fetal Weight Equations*** shown in **Figure 2** and **Supplement-Figures-II.9&10**, the average ***Error*** for predicting ***Birthweight*** among all newborns varies from as little as 18 grams for the ***Abdominal Circumference Binary Cellular Estimated Fetal Weight Equation*** (#14c) to as much 194 grams for the ***Hadlock-5 Estimated Fetal Weight Equation***. In contrast, the values for the ***Standard Deviation*** show us that the prediction of ***Birthweight*** for ∼1-in-300 fetuses is off by 892 grams, when estimated by the ***Hadlock-4 Estimated Fetal Weight Equation***, and off by 1099 grams, when estimated by the ***Hadlock-5 Estimated Fetal Weight Equation***, and off by 1029 grams, when estimated by the ***Abdominal Circumference Binary Cellular Estimated Fetal Weight Equation***. The prediction of ***Birthweight*** for ∼1-in-20 fetuses is off by 595 grams, when estimated by the ***Hadlock-4 Estimated Fetal Weight Equation***, and off by 733 grams, when estimated by the ***Hadlock-5 Estimated Fetal Weight Equation***, and off by 686 grams, when estimated by the ***Abdominal Circumference Binary Cellular Estimated Fetal Weight Equation***. The prediction of ***Birthweight*** for ∼1-in-3 fetuses is off by 297 grams, when estimated by the ***Hadlock-4 Estimated Fetal Weight Equation***, and off by 366 grams, when estimated by the ***Hadlock-5 Estimated Fetal Weight Equation***, and off by 343 grams, when estimated by the ***Abdominal Circumference Binary Cellular Estimated Fetal Weight Equation***.

##### The Source of *Estimated Fetal Weight Error* and *Standard Deviation*

As we shall see below, the source of these disappointing ***Standard Deviations*** seen for all 7 ***Estimated Fetal Weight Equations*** lies not in their calculations of ***Fetal Weight***, but in the measurement of ***Fetal Size*** by ultrasound, when the patient was examined by the sonographer. Fortunately, as we shall see, this is a problem for which ***Binary Cellular Analysis*** can offer assistance.

##### Assessment of Small and Large Newborns

Many ***Estimated Fetal Weight Equations*** are degraded by their inability to accurately capture the sizes of fetuses that will become ***Large*** and ***Small Babies***^.27,31^,***36 The INTERGROWTH*** and ***Hadlock-1,-2,-3,-4***, and ***-5 Estimated Fetal Weight Equations*** all overestimate the ***Weight*** for fetuses that become newborns of ***Small Size***, but this is not the case for the ***Abdominal Circumference Binary Cellular Estimated Fetal Weight Equation*** (#14c), which was equally reliable for fetuses that become newborns of all sizes (**Figure 3** and **Supplement-Figure-II.12**).

**Figure 3.**
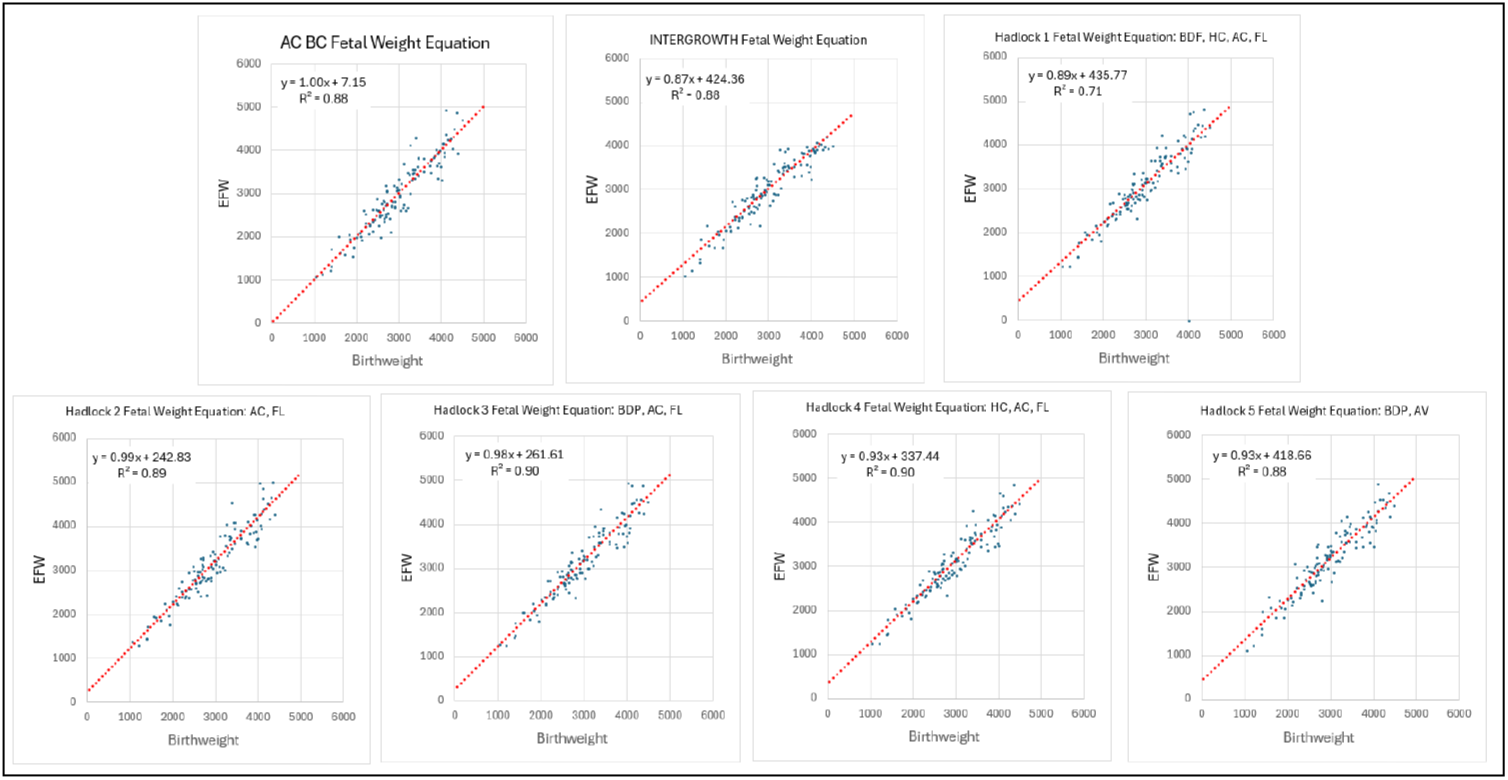
Comparison of ***Estimated Fetal Weight*** vs ***Birthweight*** for seven ***Estimated Fetal Weight Equations: Abdominal Circumference Binary Cellular Estimated Fetal Weight Equation, INTERGROWTH Estimated Fetal Weight Equation, Hadlock-1,-2,-3,-4***, and ***-5 Estimated Fetal Weight Equations***, in the ***Testing Dataset***, based on ultrasound carried out within 7 days of birth.

##### *Binary Cellular Estimated Fetal Weight Equations* of Various *Body Parts*, Individually and Averaged

The ***Abdominal Circumference Binary Cellular Estimated Fetal Weight Equation*** (#14c) proved superior to the ***Binary Cellular Estimated Fetal Weight Equations*** based on the ***Biparietal Diameter*** (BPD); ***Occipitofrontal Diameter*** (OFD); ***Femur Length*** (FL); and ***Head Circumference*** (HC) (**Supplement-Figures-II.9-18**). Despite the striking superiority of ***Abdominal Circumference Binary Cellular Estimated Fetal Weight Equation*** (#14c) to the ***Femur Length Binary Cellular Estimated Fetal Weight Equation***, and ***Head Circumference Binary Cellular Estimated Fetal Weight Equation***, averaging them yielded almost the same outcome as the ***Abdominal Circumference Binary Cellular Estimated Fetal Weight Equation*** (#14c) alone (**Supplement-Figure-II.19-23**). This suggests practical improvements of these measurements may make averaging a way to smooth out and improve the performance of any single measurement.

##### Sonographer Performance

Variability in the outcome value for a ***Fetal Ultrasound Measurements*** can be expected to come from a variety of sources, among which are: 1) Which equation is employed to interpret the ***Fetal Ultrasound Measurement***? 2) Which person is engaged in carrying out the ***Fetal Ultrasound Measurement***? An examination of results reported for 13 studies reveals an enormous amount of variation in both the ***Mean Error***, and the ***Standard Deviation*** of the ***Error*** (**Supplement-Figures-II.26 &27**), when examining the performance of a single equation, the ***Hadlock-4, Estimated Fetal Weight Equation***^21-33^. ***There is no evident correlation between Mean Error*** and ***Standard Deviation*** of the ***Error*** (**Figures 4** and **5** and **Supplement-Figure-II.26**). These studies have been carried out over four decades, but there was no indication of improvement in performance over time (**Figure 5, Supplement Figure-II.27**)

**Figure 4.**
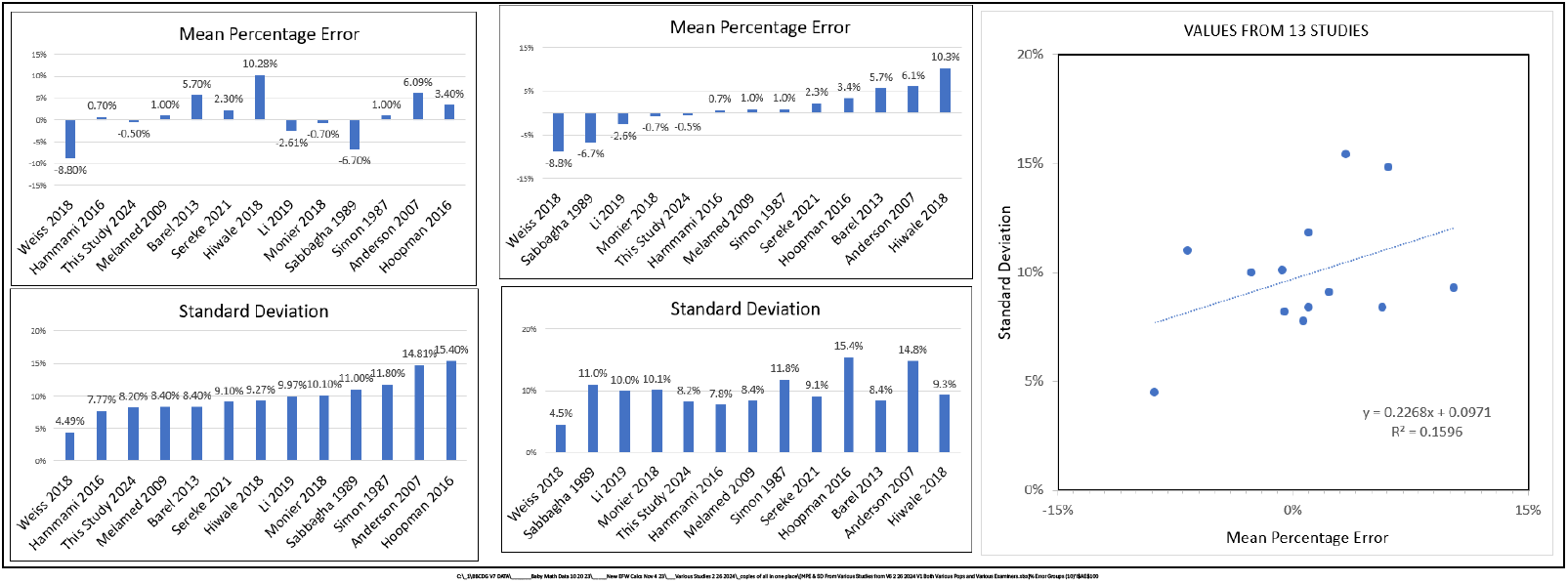
Comparison of the performance of the ***Hadlock-4, Estimated Fetal Weight Equation*** from 13 different studies.^21-33^

**Figure 5.**
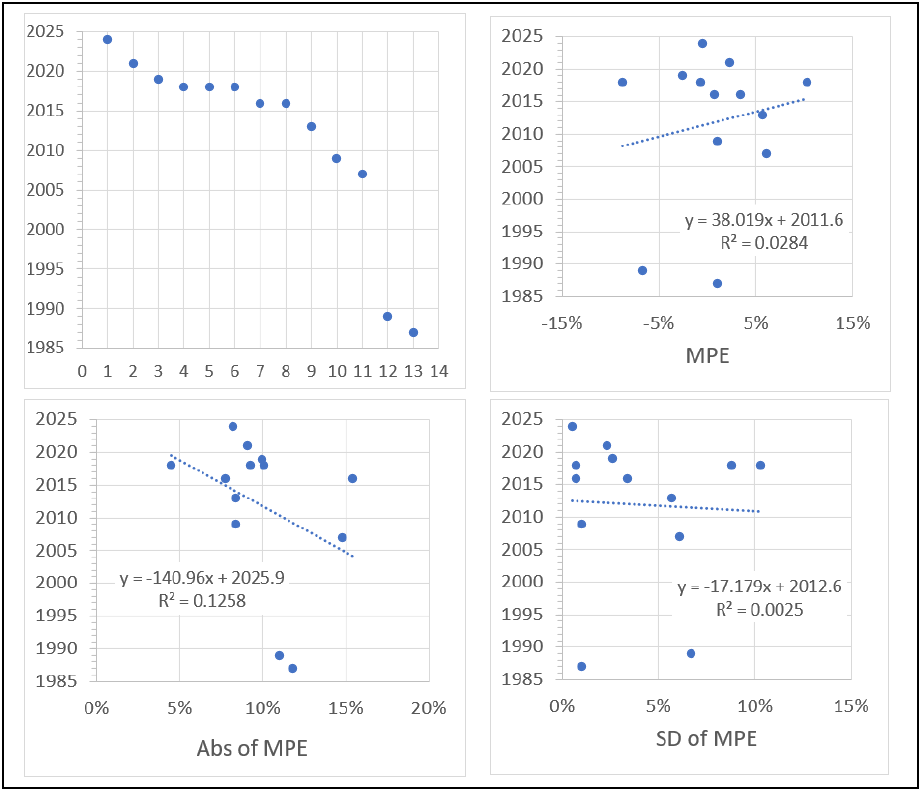
Comparison of the performance of the ***Hadlock-4, Estimated Fetal Weight Equation*** from 13 different studies, by date of study.^21-33^

A number of studies have compared the performance of individuals in carrying out fetal ultrasound. One study examined the performance of 11 sonographers, reported in terms of ***Mean Error*** and the ***Standard Deviation*** of the ***Error***.^34^ The variation from highest to lowest values was striking; the ***Mean Error*** of the ***Hadlock-4, Estimated Fetal Weight Equation*** value varied from -0.9% to -6.3%, while the ***Standard Deviation*** of the ***Error*** varied from 8.4% to 12.0% (**Figure 6** and **Supplement-Figure-II.27**). There was no evident correlation among the results of the 11 sonographers between the ***Mean Error*** and the ***Standard Deviation*** of the ***Error***.

**Figure 6.**
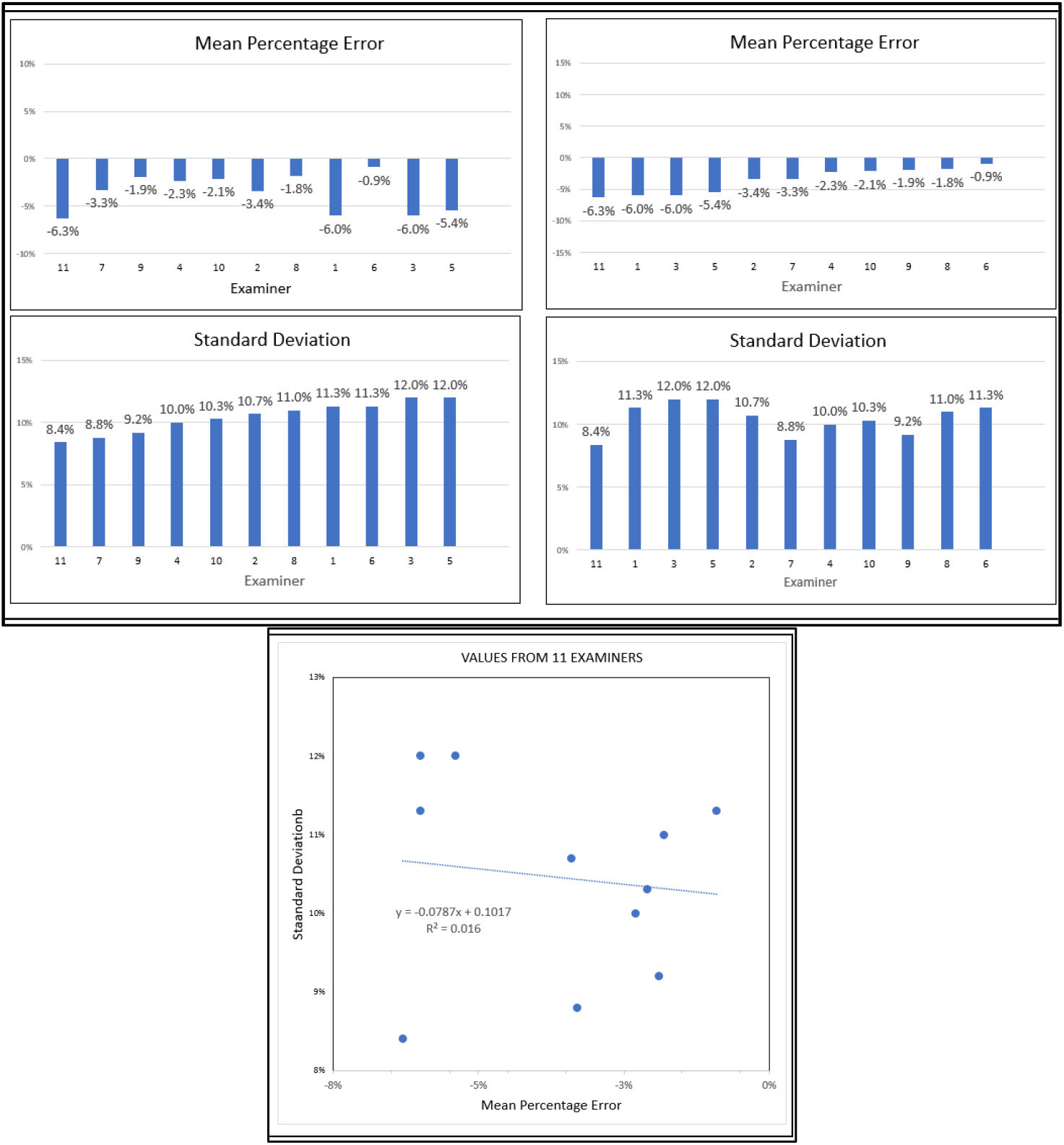
Comparison of the performance of the ***Hadlock-4, Estimated Fetal Weight Equation*** by 11 different examiners, as reported by Siemer et al^34^

Another study compared the performance of 259 ***Experienced Examiners*** with 1682 ***Inexperienced Examiners***, whose ***Ultrasound Measurements*** were applied to 11 ***Estimated Fetal Weight Equations***. The variation in performance betwewen the ***Experienced Examiners*** and ***Inexperienced Examiners*** was much greater than the performace of the various ***Estimated Fetal Weight Equations***. The correlation coeffficient (***r***^***2***^) varied by 16.7% between the ***Experienced Examiners*** and ***Inexperienced Examiners***. The ***Estimated Fetal Weight Equations*** varied much less than the ***Examiners***, ranging highest to lowest varied by 3.5% among the data collected by the the ***Experienced Examiners***, and 9.3% among the deata collected by ***Inexperienced Examiners*** (**Supplement-Figure-II.29**).

Another example of variation from site to site in the performance of fetal ultrasound could be seen by the correlation coefficient (***r***^***2***^) when comparing the results derived by the ***Hadlock-4 Estimated Fetal Weight Equation***. In the study results reported here, ***r***^***2***^=0.90, while in the study reported by Sharma et al^35^, ***r***^***2***^=0.3468 (**Supplement-Figure-II.30**).

That performance can be traced to image quality was shown by Townsend et al,^36^ who separated ***Ultrasound Measurements*** whose images they judge to be “Good” from those they judged to be “Poor”. The ***Mean Error*** of the ***Hadlock-4, Estimated Fetal Weight Equation*** value was 2.9% for studies with “Poor Images”, and twice as high, at 6%, for studies with “Good Images”. The ***Standard Deviation*** of the ***Error*** of the ***Hadlock-4, Estimated Fetal Weight Equation*** value was 8.9% for studies with “Poor Images”, and 15%, for studies with “Good Images” (**Supplement-Figure-II.31**).

***Binary Cellular*** Estimation of ***Fetal Weight*** from ***CRL*** Measurements Early in Pregnancy

To develop a method for estimating ***Fetal Weight***, early in pregnancy, we employed the ***Binary Cellular Universal Growth Equation*** (#2) and the ***CRL from Fetal Age Equation*** (#7b) to construct the ***Crown to Rump Length Binary Cellular Fetal Weight Equation***:

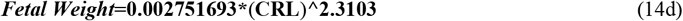

For a detailed description of the steps involved, see the SUPPLEMENT.

## DISCUSSION

### *Binary Cellular Mathematics* Provides *Estimated Fetal Weight Equations* That Are Better

Here we have seen that the ***Binary Cellular Estimated Fetal Weight Equation***, and specifically the ***Abdominal Circumference Binary Cellular Estimated Fetal Weight Equation***, performs better than six other ***Estimated Fetal Weight Equations***: the ***INTERGROWTH*** and ***Hadlock-1,-2,-3,-4***, and ***-5 Estimated Fetal Weight Equations***^.1-14^ ***The Hadlock-4 Estimated Fetal Weight Equation*** is of special note, as it is among the most widely used, and has often been found to be the most accurate and precise of the many ***Estimated Fetal Weight Equations*** developed over the past five decades.^7-14^

The ***Abdominal Circumference Binary Cellular Estimated Fetal Weight Equation*** yielded a lower ***Mean Error*** than the six other ***Estimated Fetal Weight Equations***. Furthermore, unlike these six current ***Estimated Fetal Weight Equations***, the ***Abdominal Circumference Binary Cellular Estimated Fetal Weight Equation*** showed an equal performance among fetuses that become newborns of all sizes.

The ***Hadlock-4 Estimated Fetal Weight Equation*** has a marginally lower ***Standard Deviation*** of the ***Error*** than the ***Abdominal Circumference Binary Cellular Estimated Fetal Weight Equation***, but the difference was negligible. For 2-out-of 3 ultrasound measurements, the difference between the two equations is less than 28 grams, ∼1% of the weight of an average newborn, while for 299-out-of-300 ultrasound measurements, the difference between to the equations is less than 137 grams, ∼4% of the weight of an average newborn.

### *Binary Cellular Mathematics* Makes Possible Better *Sonographer Performance* of *Fetal Weigh Estimation*

Here we have seen that the main source of error in estimating ***Fetal Weight*** lies not in the choice of the equation, but in the quality of the ***Ultrasound*** measurement carried out by the sonographer^.37,38^,39 ***Mean Errors*** and ***Standard Deviation*** of the ***Error*** seen from different institutions, and from different sonographers, were greater than the ***Mean Errors*** and ***Standard Deviation*** of the ***Error*** from different equations.^21-36^

Why don’t all sonographers perform as well as the best sonographers? One likely source of this shortcoming may well be the inability of ***Estimated Fetal Weight Equations*** to provide sonographers and their supervisors with feedback on performance, since they rely on multiple ***Fetal Ultrasound Measurements***. Even if the relationship between an ***Ultrasound Measurement*** and the subsequent ***Birthweight*** does not match up, the sonographer, and the sonographer’s supervisor, are unaware of this, because multiple ***Ultrasound Measurement*** are combined into the estimate of ***Fetal Weight. Binary Cellular Mathematics*** now makes such feedback possible, as each ***Binary Cellular Estimated Fetal Weight Equation*** relies on a single ***Fetal Ultrasound Measurement***, related to a single ***Birthweight***. Furthermore, it is impossible to “fudge” such a test, since the ***Birthweight*** is not known at the time of the ***Ultrasound Measurement***. Thus, performance feedback is now made possible with each ***Binary Cellular Estimated Fetal Weight Equation***, as each relates individual ***Ultrasound Measurement*** to individual ***Birthweights***.

### *Binary Cellular Biology* Tells Us Why Some *Ultrasound Measures* Are Better Than Others

Why does ***Abdominal Circumference*** provide a better estimate of ***Fetal Weight*** than ***Femur Length*** and ***Head Circumference***? As we outlined in the previous paper in this series, the underlying biology for why some parts of the body increase in size more rapidly than other parts of the body emerges from ***Binary Cellular Analysis*** of the growth of the ***tissues, organs***, and ***anatomical structures*** of the body. This ***Binary Cellular Analysis*** revealed that ***Body Part*** creation and growth is captured by a simple expression, the ***Binary Cellular Allometric Growth Equation*** (#11), governed by two parameters: the ***Binary Cellular Allometric Birth, B***_***N1***_ and the ***Binary Cellular Allometric Slope, S***_***N***._ The ***Binary Cellular Allometric Slope, S***_***N***_, is determined by the average ***Cell Cycle Time, c***_***p***_, of the cells in a ***Body Part***, which is specified in ***Cell-Heritable*** fashion, by a mechanism such as DNA methylation, in the part’s ***Founder Cell*** at its ***Binary Cellular Allometric Birth, B***_***N1***_. The greater the change in the ***Cell Cycle Time, c***_***p***_, of the cells in a ***Body Part***, the greater is the growth of the ***Body Part*** relative to the body as a whole, and the stronger will be the measure of the growth of the whole body from the ultrasound measurement of that ***Body Part***. Therefore, ***Binary Cellular Analysis*** shows us that it is ***Differential Cellular Proliferation***, that is, ***Cellular Selection***^,18,19^ ***that make Ultrasound Measurements*** of some ***Body Parts*** more effective for estimating ***Fetal Weight*** than other ***Body Parts***.

It is from this biology that we can understand why ***Abdominal Circumference*** has given better estimates of ***Fetal Weight*** than ***Biparietal Diameter, Occipitofrontal Diameter, Femur Length***, and ***Head Circumference***. It is also from this ***Binary Cellular Analysis*** that we can revisit ***Body Part*** characteristics, biochemically, anatomically, and morphometrically, so as to identify which ***Ultrasound Measurements*** can give the most reliable measures of ***Fetal Weight***, as we shall examine below.

### *Binary Cellular Biology* Makes it Possible to Find Better Things to Measure

Note how our ***Cellular Allometric Analysis*** of human ***Body Parts*** shown in **Supplement-Figure-I.30** and **31** shows components of the abdomen (kidney and spleen) having steeper ***Binary Cellular Allometric Slopes, S***_***N***_, than the brain. This is why the ***Abdomen*** gives a better result in predicting ***Birthweight*** than the ***Head***. It follows that revisiting ***Cellular Allometric Analysis*** of ***Body Parts***, such as that shown in **Supplement-Figure-I.30** and **31** can give insight into which ***Ultrasound Measurements*** can be expected to give the most accurate and precise measurements of ***Fetal Weight***. This also means that we can go back to the basic anatomy of human fetuses, available to us from fetal MRI data,^40^ 3D Sonography^,41,42^ or fetal autopsy imaging, by methods such as high-resolution X-ray based Micro CT^,43,44^ to search systematically for which fetal measurement should be expected to give us the most accurate and precise ***Estimated Fetal Weights. Binary Cellular Mathematics*** thus gives us the power to re-invent ***Fetal Weight Estimation*** by returning to the 3D anatomy of the human fetus with tools not available to the pioneers of this field.

### *Binary Cellular Mathematics* Can Combine *Ultrasound Measurements* of Multiple Anatomical Sites

Here we have seen that ultrasound measurements of multiple anatomical sites can be combined by utilizing the ***Binary Cellular Estimated Fetal Weight Equation*** for each anatomical site, and then averaging the estimates into a single value for ***Fetal Weight***. Measurement of performance by the sonographer is thus retained. Furthermore, as we shall see in the next paper in this series,^15^ ***Binary Cellular Analysis*** provides a basis by which multiple measurements made at different points in time can also be combined.

### Web-based Calculators Will Provide Clinicians with *Binary Cellular* Calculations That Aid Obstetric Care

A first-generation Web-based calculator for calculating ***Fetal Weight*** from the ***Abdominal Circumference Binary Cellular Estimated Fetal Weight Equation***, is now available at “https://kidzgrowth.com“. Encoding the additional ***Binary Cellular*** calculations described here is straightforward,^45^ and underway. Also to be provided by these calculators will be tools for performance feedback, comparing each ***Ultrasound Measurement*** to the ultimate ***Birthweight***, so that sonographers and their supervisors can review performance over the long term, as well as the performance of each individual case. These web-based calculators will make the results of these ***Binary Cellular*** mathematical manipulations widely available for aiding in obstetric care of individual pregnancies.

## Supporting information

SUPPLEMENT Data and Details

